# The systemic anti-diabetic effect of polyherbal formulation *Varanadi Kashayam* is mediated through GLP-1 secretion and DPP4 inhibition

**DOI:** 10.1101/2024.06.29.601306

**Authors:** Anjana Thottappillil, Sania Kouser, Abhi V. Badiger, Priyanka Gladys Pinto, Srimathy Ramachandran, Suresh Janadri, Manjunatha P. Mudagal, Subrahmanya Kumar Kukkupuni, Suma Mohan S, Chethala N. Vishnuprasad

## Abstract

**Background:** Frontiers of health science increasingly emphasizes systems and network medicine approaches for managing complex lifestyle diseases such as diabetes. In this context, systemic targets like incretin hormones and their modulators - particularly DPP4 inhibitors - have gained prominence. *Ayurveda*, the Indian System of Medicine (ISM), with its clinically validated multicomponent formulations offers a valuable resource for integrative therapeutic strategies with systemic mode of action.

**Purpose:** The study investigates the incretin modulatory effect of *Varanadi Kashayam* (VA), an *Ayurveda* polyherbal formulation used in the clinical management of diabetes and its comorbidities.

**Experimental approach:** A high-fat diet induced Sprague-Dawley (SD) rat model was used to study the anti-diabetic, GLP-1 secretory, and anti-obesity effects of VA. *In vitro* studies using GLUTag cells assessed the GLP-1 secretion and DPP4 gene expression modulation; and studies using 3T3-L1 fibroblasts examined the anti-adipogenic effects. Computational methods including molecular docking and molecular dynamic simulations were used to identify the phytochemicals responsible for DPP4 inhibition.

**Results:** VA administration significantly improved fasting blood glucose and oral glucose tolerance in experimental animals, along with enhanced GLP-1 secretion. The *in vitro* results showed inhibition of DPP4 enzyme activity, increase in GLP-1 secretion and downregulation of DPP4 gene expression in GLUTag cells, and suppression of adipogenesis in 3T3-L1 fibroblast cells. Computational analyses ascertained Chebulinic acid, Chebulagic acid and Terchebin as top ranked phytochemicals responsible for DPP4 inhibition effect.

**Conclusion:** Combining *in vitro*, *in vivo* and *in silico* findings, this study provides valuable insight into the incretin modulatory effect of VA in treating diabetes and associated metabolic diseases.

**Graphical Abstract:** 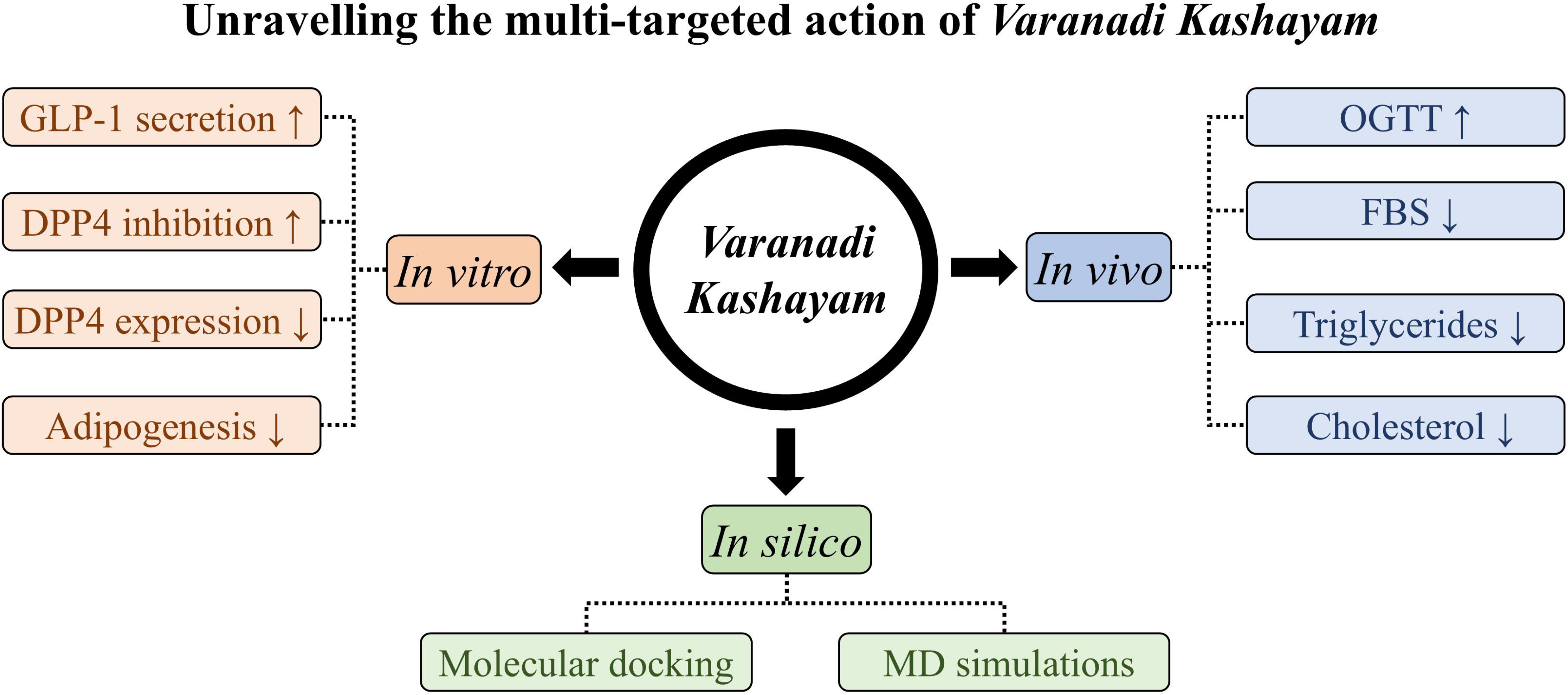

**Highlights:**

- *Varanadi Kashayam* (VA) improved blood glucose and GLP-1 levels in diabetic rats.
- Inhibited DPP4 activity and gene expression in GLUTag cells.
- Suppressed adipogenesis and adipogenic markers in 3T3-L1 cells.
- Chebulinic acid, Chebulagic acid, and Terchebin identified as key DPP4 inhibitors.

## 1. Introduction

Incretin-based therapies - such as glucagon-like peptide-1 (GLP-1) analogues, GLP-1 secretagogues, and GLP-1 receptor agonists as well as dipeptidyl peptidase-4 (DPP-4) inhibitors - have emerged as promising strategies for the treatment of type 2 diabetes (T2D).^1,2^ GLP-1, the key incretin hormone produced by enteroendocrine cells, plays a central role in maintaining the body’s glucose homeostasis by enhancing nutrient- and glucose-mediated insulin secretion from pancreatic beta cells, suppressing glucagon release, and slowing gastric emptying. It is also known to exert its biological action on other tissues of the body, thus regulating diverse biological functions.^3^ However, GLP-1 is rapidly degraded by the enzyme DPP-4, a serine protease inhibitor whose concentration and activity is often elevated in diabetic individuals, and thereby reduces the incretin effect.^4^ Given the multi-targeted and systemic action of GLP-1 in glucose metabolism, there is growing interest in deepening the understanding of GLP-1 biology and identifying novel analogues and secretagogues of GLP-1 as well as inhibitors of the DPP4 enzyme.^5,6^ These strategies are expected to improve the incretin effect of the body, which in turn helps in reducing both fasting and postprandial blood glucose levels.

Diabetes is perhaps one of the most complex metabolic diseases associated with various comorbidities like obesity and macro- and micro-vascular complications. Managing T2D and restoring glycemic control is not only essential for reducing hyperglycemia, but also for preventing a myriad of comorbidities associated with diabetes. Unfortunately, the current drug therapies are addressing only the hyperglycemia and often fail to adequately address the systemic biological changes and comorbidities associated with these conditions.^7^ Therefore, it is imperative to formulate new concepts and strategies that can address the systems biology of glucose metabolism to effectively manage T2D and associated comorbidities.

*Ayurveda*, the ancient Indian system of medicine, provides holistic approaches to managing complex diseases like diabetes and obesity. Central to the doctrine of *Ayurveda* is to secure the physiological and psychological homeostasis of the body for health and disease management. By integrating ayurvedic principles into modern healthcare, we can develop novel strategies that address both the symptomatic and systemic biology of disease to provide a more comprehensive and effective management. The concept of *Agni* (the digestive fire) encompasses gastrointestinal and cellular metabolism that governs the transformation of food into energy and nutrients and is considered a key factor in maintaining the body’s metabolic homeostasis.^8^ When *Agni* is in balance, it ensures optimal digestion, absorption, and assimilation of nutrients, promoting overall health and well-being. In *Ayurveda*, maintaining balanced *Agni* is considered crucial for preventing and managing metabolic disorders. Consequently the dietary guidelines, lifestyle modifications, and therapeutic interventions are primarily directed towards supporting and managing *Agni* functions.^8^

Integrating the concepts of incretin biology and Ayurvedic principles of systemic diabetes management, the current study explores the incretin modulatory effect of *Varanadi Kashayam* (VA), a polyherbal formulation prescribed for the clinical management of diabetes and various other predisposing factors and comorbidities of diabetes like obesity, insulin resistance, NAFLD etc.^9^ VA consists of 16 medicinal herbs (detailed in Table 1), several of which have been documented for their anti-diabetic and hypoglycemic effects.^10^ VA, as per the classical *Ayurveda* text *Ashtanga hrudaya*, is indicated for improving sluggish metabolism and reducing *meda* or fat.^11^ From an *Ayurveda* perspective, one of the primary pharmacological actions of VA is modulation of *Agni*, both at gastro-intestinal tract (*Jatharagni*) and at peripheral tissues (*Dhatwagni*).^12^ Owing to its GIT centred pharmacological action in the context of diabetes and obesity, our study explores the incretin modulatory effects of VA using integrated approach that includes *in vitro* assays*, in vivo* animal models and *in silico* computational analysis.

**Table 1:**
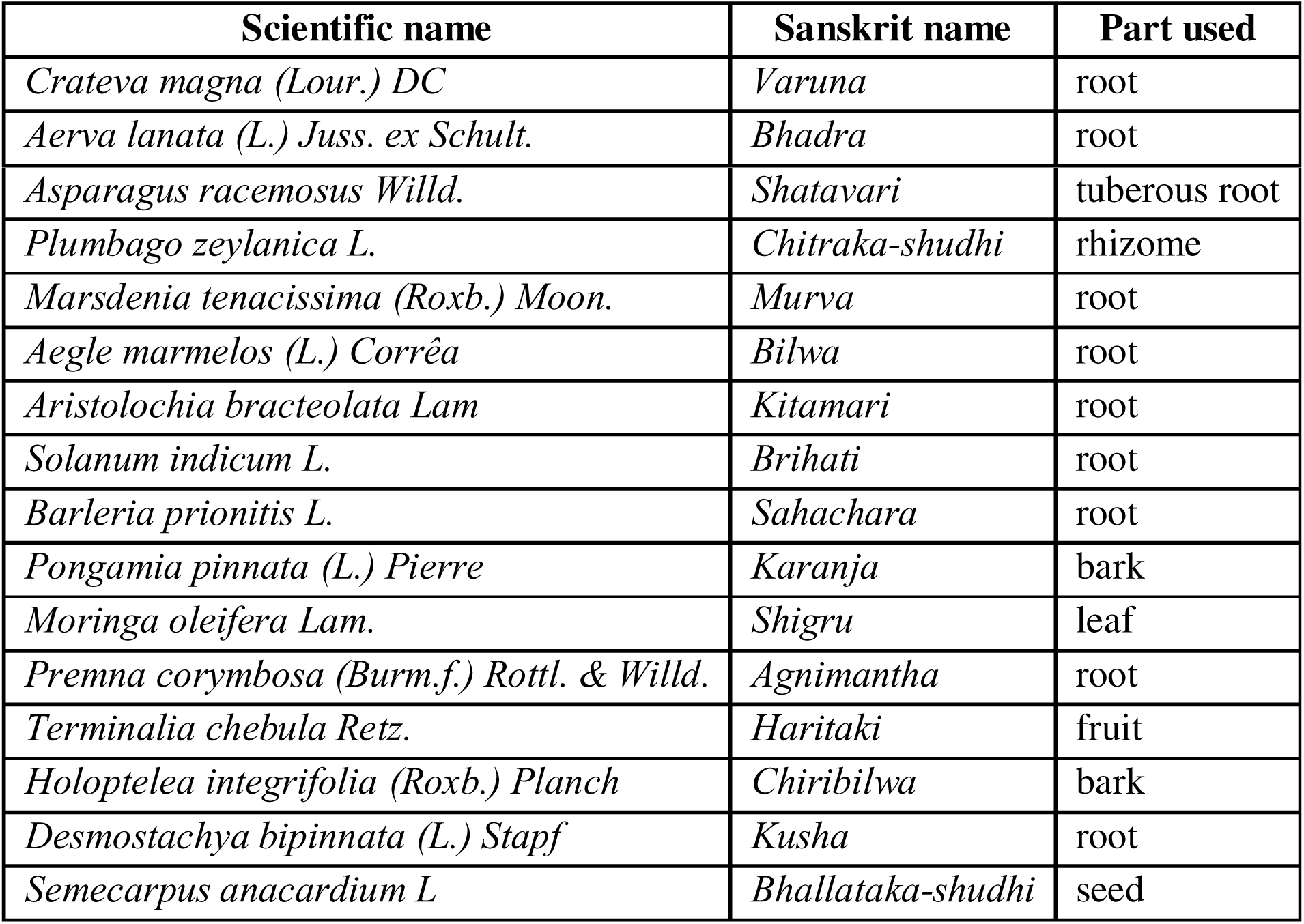
Table shows the list of plants present in VA along with their Sanskrit name and the parts used in the drug preparation.

## 2. Materials and Methods

### 2.1 Chemicals and Reagents

VA was procured from a leading *Ayurveda* drug manufacturer (The Arya Vaidya Pharmacy, Tamil Nadu with batch no: AEA156 (for *in vitro* studies) and batch no: AII-092 (for *in vivo* studies). The whole formulation was used in its original form without any processing or extraction. Fine chemicals were purchased from Sigma-Aldrich (St. Louis, MI). Dulbecco’s Modified Eagle’s Medium (Cat. No. 11965084) and Foetal Bovine Serum (Cat. No. 10270106) were purchased from Thermo Fisher Scientific Inc. The DPP4 drug discovery kit was procured from Enzo Life Sciences (Farmingdale, New York, NY, USA). The GLP-1 kit was purchased from Ray biotech (CAT NO - EIAR-GLP 1-1, RAT GLP-1 EIA,). Triglyceride and Cholesterol estimation kits were purchased from Biosystems (Cat. No. 11258 and 11505 respectively). Glycomet (Metformin 500 mg Tablet) and Metformin hydrochloride (Honeychem pharma) were purchased from authentic sources.

For quantitatively using VA formulation for various *in vitro* assays, total tannins present in the formulation was estimated using the standard Folin - Ciocalteu method using gallic acid as standard.^13^ Briefly, 10 μL of the formulation mixed with 40 μL of water, 50 μL Folin’s reagent and 100 μL of 3.5% Na_2_CO_3_ was incubated at room temperature for 30 min. A set of gallic acid standards (50, 25, 12.5, 6.25, 3.125 μg/mL) were prepared in the same manner. The absorbance was measured at 700 nm using a multi-well plate reader (xMark Microplate Spectrophotometer, BioRad, USA). The experimental concentrations of test samples were expressed as ‘μg of gallic acid equivalent tannin (GAE)/mL of sample.

### 2.2 Dipeptidyl peptidase-4 (DPP4) inhibition assay

The DPP4 inhibition assay was carried out as per the kit instructions. Briefly, various concentrations of VA were prepared with the assay buffer. The test sample (VA), positive control and the standard inhibitor (given in the kit) were incubated with DPP4 enzyme for 20 minutes at room temperature. The fluorogenic substrate provided with the kit was added and the plate was read for 30 mins using a fluorometer (Biotek Synergy H1 microplate reader) at Ex:380/EM:460 nm. The percentage remaining activity in the presence of inhibitor was calculated using the formula below

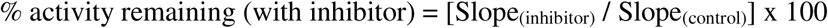

### 2.3 GLP1 secretion and DPP4 expression studies

Murine GLUTag cell line was obtained as a generous gift from Prof. Tohru Hira, Hokkaido University, Japan with the kind permission of Prof. Daniel Drucker, University of Toronto, Canada. The cells were maintained in DMEM containing 1% penicillin and streptomycin supplemented with 18% FBS, at 37°C in a 5% CO_2_ humidified atmosphere. For the assays cells were seeded in 24-well plates (85000 cells/well) and grown for 48 h and then serum starved for one hour. Following which, the cells were treated with glucose free media (DMEM) without or with different concentrations of VA (12.5, 25 and 50μg of GAE/mL) and incubated for 2 h in a 5% CO_2_ atmosphere at 37°C. The media was collected and briefly centrifuged to remove any debris. The secreted GLP-1 levels were assayed using GLP-1 ELISA kit (Ray biotech) following the manufacturer’s instructions. The OD was read at 450 nm as per kit protocol.

For DPP4 expression studies, total RNA was extracted from GLUTag cells, (treated with and without VA) using Trizol reagent (RNAiso Plus, Takara). One microgram of total RNA was subjected to cDNA synthesis using PrimeScript RT Reagent kit (Takara). The target cDNA was amplified using TB Green Premix Ex Taq II (Takara) and the following sense and antisense primers: FP - TCAACAGTCATGAGCAGAGC, RP – GGTCTTCATCCGTGTACCAC.

### 2.4 Anti-adipogenic effect of VA on 3T3-L1 cells

The 3T3-L1 fibroblasts were purchased from the National Centre for Cell Sciences, Pune, India.The cells were maintained in DMEM containing 1% penicillin and streptomycin supplemented with 10% FBS at 37°C in a 5% CO_2_ humidified atmosphere. The 3T3-L1 fibroblasts were differentiated by incubating two days post-confluent (day-0) plates in differentiation induction medium containing 500μM IBMX, 250nM dexamethasone and 50nM insulin (MDI). On day-3, the medium was replaced with insulin media for 2 days and subsequently the cells were maintained in fresh culture media till they attain complete adipocyte morphology.^14^ To check the anti-adipogenic effect of VA, 2µg of GAE/mL VA was added along with MDI and incubated. On day-7 fully differentiated adipocytes were used for Oil Red O staining, triglyceride assay and qRT-PCR.

#### 2.4.1 Oil-Red-O staining

Oil Red O staining was performed after washing the cells twice with a phosphate buffer followed by fixing with 4% formaldehyde for 30 mins. Cells were then stained with freshly diluted Oil Red O solution (0.5% Oil Red O in isopropanol diluted to 3:2 with H_2_O, and filtered) for 20 mins at 37°C. Images of cells stained with Oil Red O were captured using phase contrast microscope (Olympus-IX-71, Olympus America Inc, USA). For quantitative measurement, the stain retained was eluted by 100% isopropanol and absorbance was read at 540 nm.

#### 2.4.2 Triglyceride estimation

The cellular triglyceride accumulation was measured after scraping the cells into 250μl 0.1% PBST and pulse sonicated with 45% amplitude for 45 seconds. The lysates were assayed for their total triglyceride content using assay kit (BeneSphera) and cellular protein was estimated using the BCA method. The triglyceride content was normalized to the total protein and the results were expressed as μg of triglyceride per mg of total protein.

#### 2.4.3 Gene expression studies

Total RNA was extracted from differentiated 3T3-L1 adipocytes (treated with and without VA) using Trizol reagent (RNAiso Plus, Takara). One microgram of total RNA was subjected to first strand cDNA synthesis using PrimeScript RT Reagent kit (Takara). The target cDNA was amplified using TB Green Premix Ex Taq II (Takara) and the following sense and antisense primers: *Ppar*γ (FP: TTTTCAAGGGTGCCAGTTTC, RP: AATCCTTGGCCCTCTGAGAT), *Fabp4* (FP: AAGGTGAAGAGCATCATAACCCT, RP: TCACGCCTTTCATAACACATTCC), *Srebp-1c* (FP: GTGAGCCTGACAAGCAATCA, RP: GGTGCCTACAGAGCAAGAGG), *Fasn* (FP: TGGGTTCTAGCCAGCAGAGT, RP: ACCACCAGAGACCGTTATGC), and *Gapdh* (FP: ACCCAGAAGACTGTGGATGG, RP: CACATTGGGGGTAGGAACAC). The analysis was conducted using the ΔΔCT method. The experiment was independently repeated three times.

### 2.5 *In vivo* studies using high fat and streptozotocin treated rat model

All *in-vivo* experiments were carried out in Acharya & BM Reddy college of Pharmacy, Bengaluru, India. Male SD rats weighing 150-180 g were procured and maintained in Acharya & BM Reddy college of Pharmacy animal housing facility. Animals were maintained under controlled environmental conditions (temperature 21-25°C and normal humidity conditions). The animals were fed with commercial High Fat Diet (HFD) diet procured from VRK nutritional solutions, Pune, India. The composition of HFD and the normal diet are given in Table 2. The study protocol was approved by the institutional animal ethics committee (IAEC/ABMRCP/2023-2024/5).

**Table 2.**
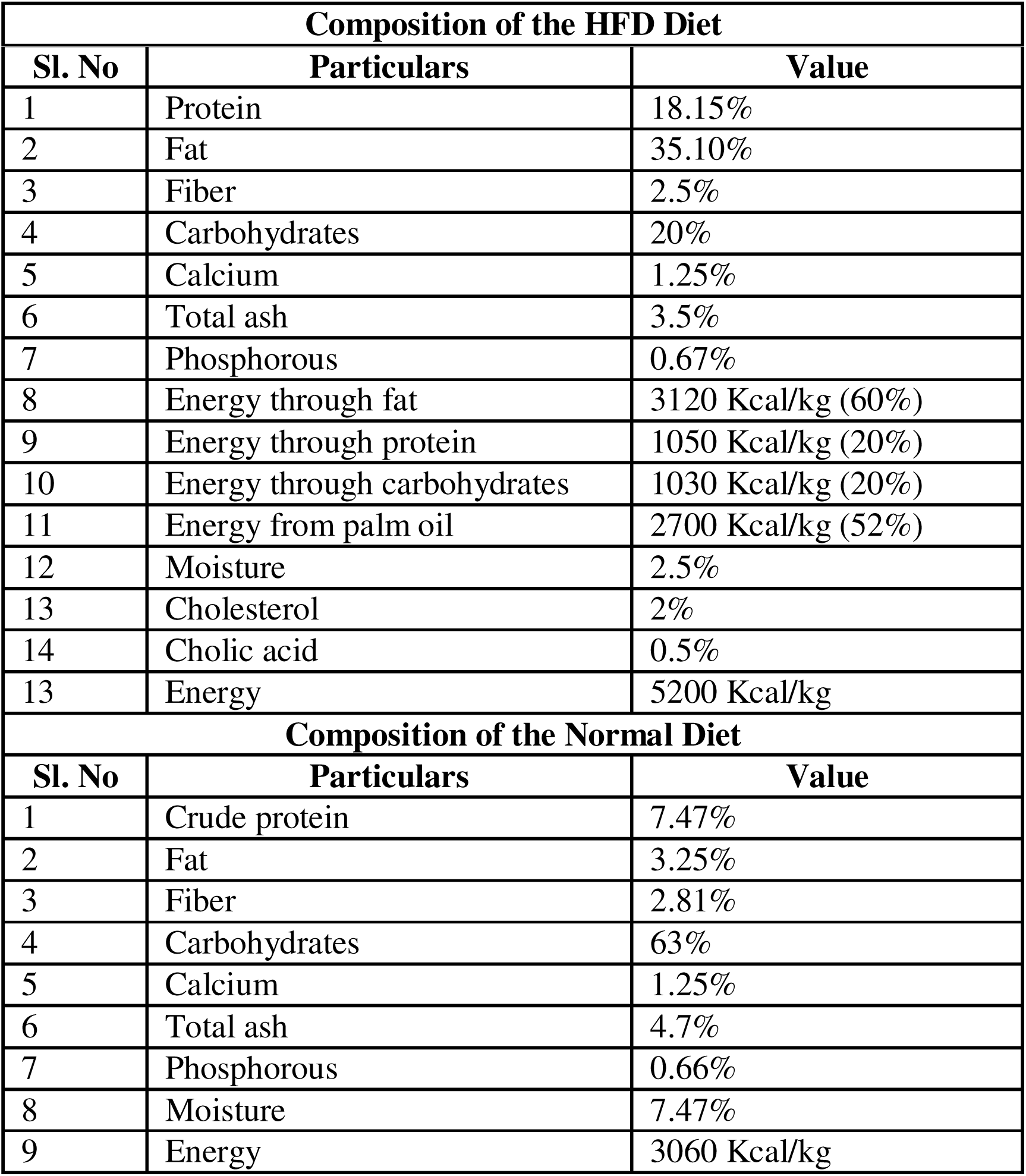
The composition of high fat and normal diet.

#### 2.5.1 Acute toxicity studies

Based on the limit test standard of the Organization for Economic Cooperation and Development (OECD) No. 425 Guideline described in the reference for acute toxicity testing of drugs in rodents, 2 female SD rats were administered orally with 2mL/Kg of VA.^15^ After 24 hrs of observation for physical and behavioural change, the animals were further administered with a higher dose of 5mL/Kg of VA and observed for 14 days for any sign of toxicity. Formulations were well tolerated and showed no fatality or no signs of any abnormal behaviour.

For administration of formulations in diabetic condition, the doses were calculated according to the conversion table based on surface area as rat equivalent volume of adult human dose, derived by multiplying the adult human dose by 0.018 per 200 g of body weight of rat.^16^ The adult human dose of formulations was considered to be 30 ml as high dose and 15 ml as low dose after discussion with the *Ayurveda* physicians. Formulation was administered by oral gavage.

#### 2.5.2 Feeding of high fat diet and induction of diabetes in experimental animals

Male SD rats weighing 150±180g were randomly divided into two nutritional groups: a standard diet (control group) and a high-fat diet (HFD - 60% energy from fat, 20% from carbohydrate, and 20% from protein; Total energy 5200 kcal/kg). Before the HFD feeding, animals were weighed and their initial blood glucose levels were recorded. The animals were fed a high fat diet and daily food intake was recorded. Animals were weighed weekly to evaluate the weight gain. After 45 days of HFD diet, animals were fasted for 3 hrs (with free access to water) and injected with streptozotocin at a dose of 30 mg/kg (STZ) prepared in citrate buffer.; pH: 4.4.^17^ The control animals received an IP injection of citrate buffer solution (vehicle). After 72 hrs of STZ injection, the fasting blood glucose was measured and rats having blood glucose level ≥ 200 mg/dL were grouped into 4 groups as shown in Table 3. The experimental rats were administered with the respective drugs with an oral gavage for a period of 30 days after STZ induction. The high fat diet was continued throughout the formulations’ treatment for the respective groups. The standard drug metformin was administered to the animal at a dose of 10 mg/200g for the first 2 weeks, and later was increased to 20 mg/200g.

**Table 3.** Table outlining the treatment conditions administered to experimental animal groups.

| Sl. No. | Group | Treatment | No. of animals |
| --- | --- | --- | --- |
| 1 | Control | Placebo | 6 |
| 2 | HFD+STZ (HFDS) | Placebo | 6 |
| 3 | Metformin | 10mg/200g BW for first two weeks<br>20mg/200g BW for last two weeks | 6 |
| 4 | VA HD | 540mg/200g BW | 6 |
| 5 | VA LD | 270mg/200g BW | 6 |

Though the animal group had six animals to begin with, due to the observed mortality with STZ induction and the resultant hyperglycemia, the results of all further experiments are expressed for the animal group containing 4 animals (n=4).

#### 2.5.3 Oral glucose tolerance test (OGTT)

The OGTT was performed after 45 days of HFD, and repeated after STZ induction on the 48th day and finally on 26^th^ day after drug intervention. Briefly, on the day of OGTT, animals were fasted for 3 hrs, following that the animals were administered with 2 gm/Kg of glucose orally. Blood samples were collected from the caudal vein, by means of a small incision at the end of the tail, at 0, 30, 60 and 120 min after glucose administration. Blood Glucose Level (BGL) was estimated by the enzymatic glucose oxidase method using a commercial glucometer (Accucheck Active glucometer and glucose strips). The results were expressed as the integrated area under the curve for glucose (AUC glucose), which was calculated using GraphPad Prism 10.1.1 Version.

#### 2.5.4 Evaluation of various biochemical parameters upon treatment with formulations

At the end of the experimental day, blood was collected from the retro-orbital sinus and centrifuged at 3500 rpm for 10 min at 4°C. The supernatant was obtained for GLP-1 measurement. The plasma samples collected from the animals were assayed for GLP-1 secretion using Raybiotech GLP-1 kit, catalogue no: EIA-GLP1. Plasma was separated and analyzed spectrophotometrically for triglyceride and cholesterol levels using the standard assay kits.

#### 2.5.5 Histopathological examination of Liver, Kidney, Pancreas and Intestine in the animal groups

At the end of the experiments, the liver, kidney, pancreas and intestine in each animal group were immediately removed and fixed in 10% formalin. The organs from a single animal representing each group were given for histological studies. The specimens were submitted to Dr Vamshi’s Biological Sciences and Research Center, Rajajinagar, Bangalore for detailed examination of the tissues.

### 2.6 Phytochemical data collection

The phytochemicals present in the 16 component plant ingredients of VA were extracted from two publicly available databases viz. Dr Duke’s Phytochemical and Ethnobotanical Database (https://phytochem.nal.usda.gov) and IMPPAT - Indian Medicinal Plants, Phytochemistry and Therapeutics (https://cb.imsc.res.in/imppat/home).^18^ Phytochemicals reported from the respective parts used in the formulation were shortlisted and considered for further studies. The detailed chemical information about the shortlisted phytochemicals were obtained from the PubChem database (https://pubchem.ncbi.nlm.nih.gov/) and used for molecular docking and other bioinformatics studies.

### 2.7 Docking of compounds to DPP4 and MD simulation

The docking of the compounds identified from VA with DPP4 using Glide was done as following the methodology reported by Thottappillil et. al. 2023 earlier from our lab.^19^ The Prime MM-GBSA (v3.0) module of the Schrodinger Maestro suite was used for binding free energy calculations of the protein-ligand complexes. Molecular dynamics simulations were performed using the Desmond (v7.3) package with an OPLS4 force field for energy calculation.^19^ The protein-ligand complexes were solvated with the TIP4P water models and simulated in an orthorhombic box with the periodic boundary conditions of size 10 × 10 × 10 Å. The system is neutralised by adjusting the number of ions and the salt concentration was maintained at 0.15M. To equally distribute the ions and solvent around the protein-ligand complex, each system was equilibrated using NPT (number of particles, system pressure and temperature) with a constant temperature of 300K. To perform interaction analysis, we used a Simulation Interaction Diagram to calculate the RMSD of the protein and ligand, ligand interactions etc. from the simulation trajectory.

### 2.8 Statistical Analysis

A one-way ANOVA test was performed to determine the significance of test samples compared to the controls and a value of p<0.05 was considered as significant.

## 3. Results

### 3.1 VA Enhances GLP-1 Secretion and Improves Diabetic Pathophysiology in HFD-STZ Induced Rats

In our study diabetes was induced in SD rats by administering a combination of HFD and STZ as outlined in Materials and Methods. An increased fasting blood glucose and impaired oral glucose tolerance confirmed the diabetic physiology in the animals. Reduction in plasma GLP-1 levels of HFD-STZ treated animals indicates a defective incretin response, which is a hallmark of chronic diabetes.^20, 21^ 30 days treatment of VA showed a concentration-dependent increase in GLP-1 level in diabetic rats (n=4) as compared to the untreated HFD-STZ group (Fig. 1A). Corroborating with this observation, the VA treated rats showed a significant reduction of fasting blood glucose level ranging from 1.5 to 1.78 fold respectively with high dose (HD - 540 mg/200 gm) and low dose (LD - 270 mg/200 gm) (Fig. 1B). Our results showed that the reduction of fasting blood glucose in high dose treatment of VA was comparable to metformin treatment whereas the low dose showed a better effect than the metformin. Additionally, VA treatment showed an improvement of glucose tolerance as evidenced by a decrease in the area under curve (AUC) of blood glucose (Fig. 1C). The slight improvement in OGTT response seen in Fig. 1C, though not statistically significant, could be due to improved incretin response. The results indicate a possible incretin modulation effect by VA, which is important in whole-body glucose homeostasis.

**Fig. 1.**
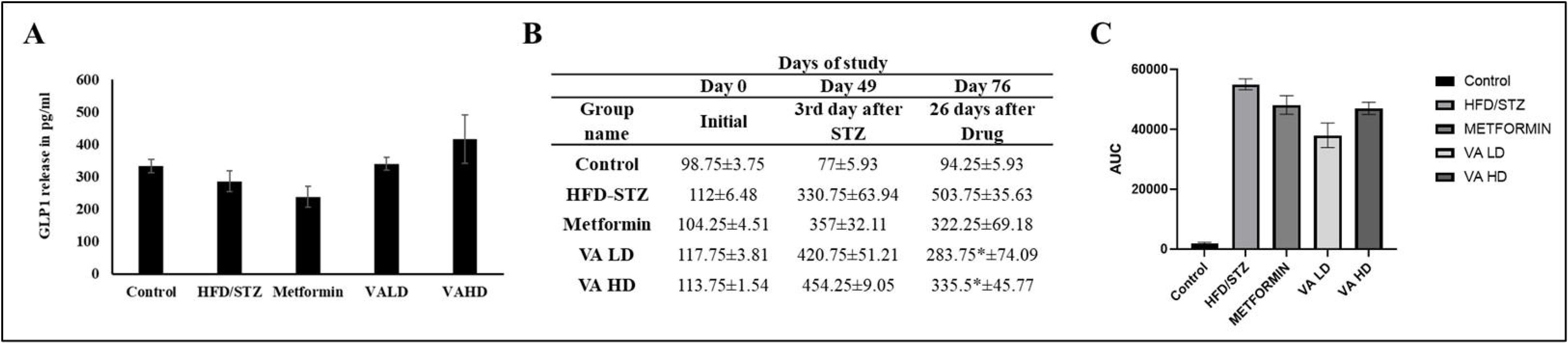
The effect of VA treatment on plasma GLP1 and Fasting Blood Glucose levels, Improvement of oral glucose tolerance in HFD-STZ rats. **(A)** Graph showing increased GLP1 levels in VA treated groups when compared to the untreated diabetic control HFD-STZ group. **(B)** Table shows the changes in fasting blood glucose (FBS) levels in experimental groups on day 0, three days after STZ treatment and 26 days after drug treatment. **(C)** Graph showing the reduction of AUC of blood glucose in VA treated groups of animals indicating an improved oral glucose tolerance in HFD-STZ animal models. Data represented as mean with standard error of mean (SEM), n=4. (p value ≤ * 0.05)

### 3.2 VA reduced body weight and plasma lipid parameters in HFD-STZ induced diabetic rats

Treatment of HFD-STZ rats with VA at both high and low doses resulted in body weight changes, as shown in Table 4. Following STZ induction, a decrease in body weight was observed across all groups, a well-documented effect of STZ administration.^22^ Fifteen days after the initiation of formulation treatment, the HD group exhibited significant weight loss, whereas no notable change was observed in the LD group. By the end of the treatment period, the reduction in body weight was no longer significant, suggesting a potential protective effect of the formulation.

**Table 4.**
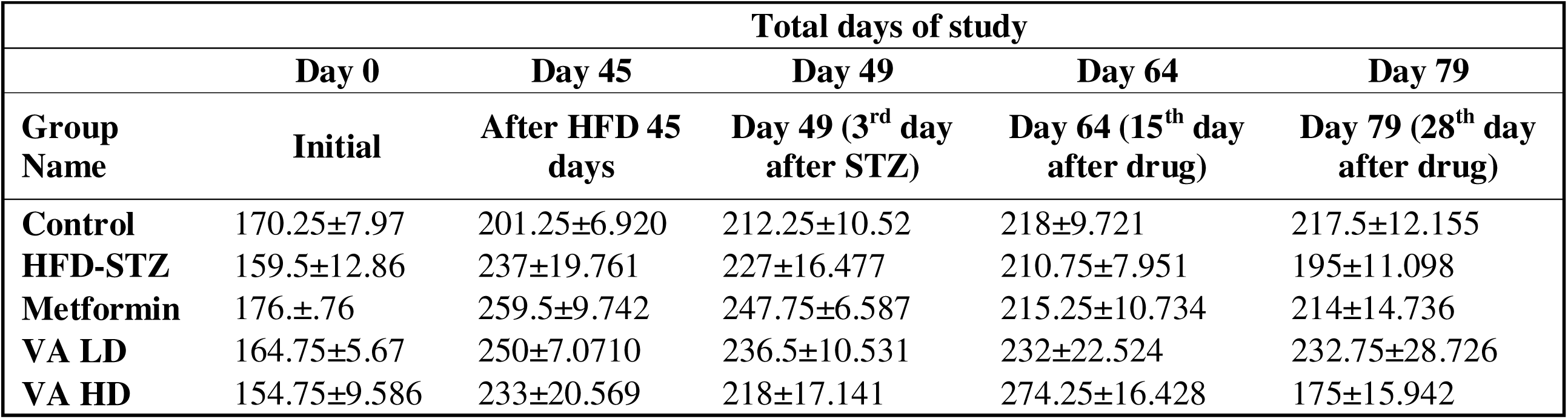
The effect of VA treatment on body weight in HFD-STZ rats. The table shows the changes in body weight at different time points, initial, after STZ induction, 15th day and 28th day after formulation treatment. Data represented as mean with SEM, n=4.

Similarly, VA treatment in HFD-STZ rats led to a decrease in plasma triglyceride and cholesterol levels in both the LD and HD groups, as presented in Table 5. These findings indicate an antihyperlipidemic effect, supporting the potential clinical application of VA as an anti-obesity treatment. Further investigation into the effects of a lower dose of VA may be beneficial to fully elucidate its mode of action, as the lower dose appeared to help maintain body weight throughout the study. These results are in line with the observations from a previous study conducted on VA by Chinchu et. al., 2020a and further validate the consistency of the biological effects of the formulation.^23^

**Table 5.** The effect of VA treatment on plasma lipid parameters in SD rats. The table shows reduction in triglycerides and cholesterol in VA treated animals when compared to the untreated HFD-STZ group. Data represented as mean with SEM, n=4.

| Group name | Control | HFD/STZ | Metformin | VALD | VAHD |
| --- | --- | --- | --- | --- | --- |
| **Triglyceride in mg/dL** | 107.7±27.7 | 215.74±52.4 | 261.76±44.9 | 171.17±20.75 | 121.85±29.55 |
| **Cholesterol in mg/dL** | 27.75 ±5.37 | 286.88±74.37 | 488.82±149.21 | 269.51±127.55 | 194.12±65.23 |

### 3.3 VA reduced adipogenesis in 3T3-L1 cells

One of the indications of VA is its *Medohara* property which can be broadly correlated to anti-adipogenic and anti-lipidemic effects. An earlier study reported by Chinchu et. al., 2020b has shown that different organic solvent extracts of VA inhibit adipogenesis in 3T3-L1 adipocytes through downregulation of key genes like *Ppar-*γ*2*, *C/ebp-*α, *Fas*, *Ap2* and *Lpl*.^24^ In our current study also, the 3T3-L1 cell model was treated with various concentrations of VA in its original form as opposed to the organic solvent extract used in the earlier study. Our results of Oil Red O staining (Fig. 2A-C) showed that accumulation of lipid droplets within the cells increased 2.35 fold upon induction with MDI, whereas co-treatment of MDI with 2µg of GAE/mL of VA significantly reduced lipid droplets accumulation compared to MDI treatment (Fig. 2D). Similar changes are observed with triglyceride levels as well (Fig. 2E). Further, similar to the earlier study, the molecular level gene expression analysis revealed that treatment with VA significantly downregulated key adipogenic markers, *Ppar*γ, *Srebp-1c*, *Fabp4* and *Fasn* by 117.05-fold, 3.14-fold, 321.07-fold and 5.19-fold respectively, when compared to MDI treated cells (Fig. 2F-I). These results, together with animal studies, support the *Medohara* property of VA and provide a molecular-level explanation for its *Medohara* action.

**Fig. 2.**
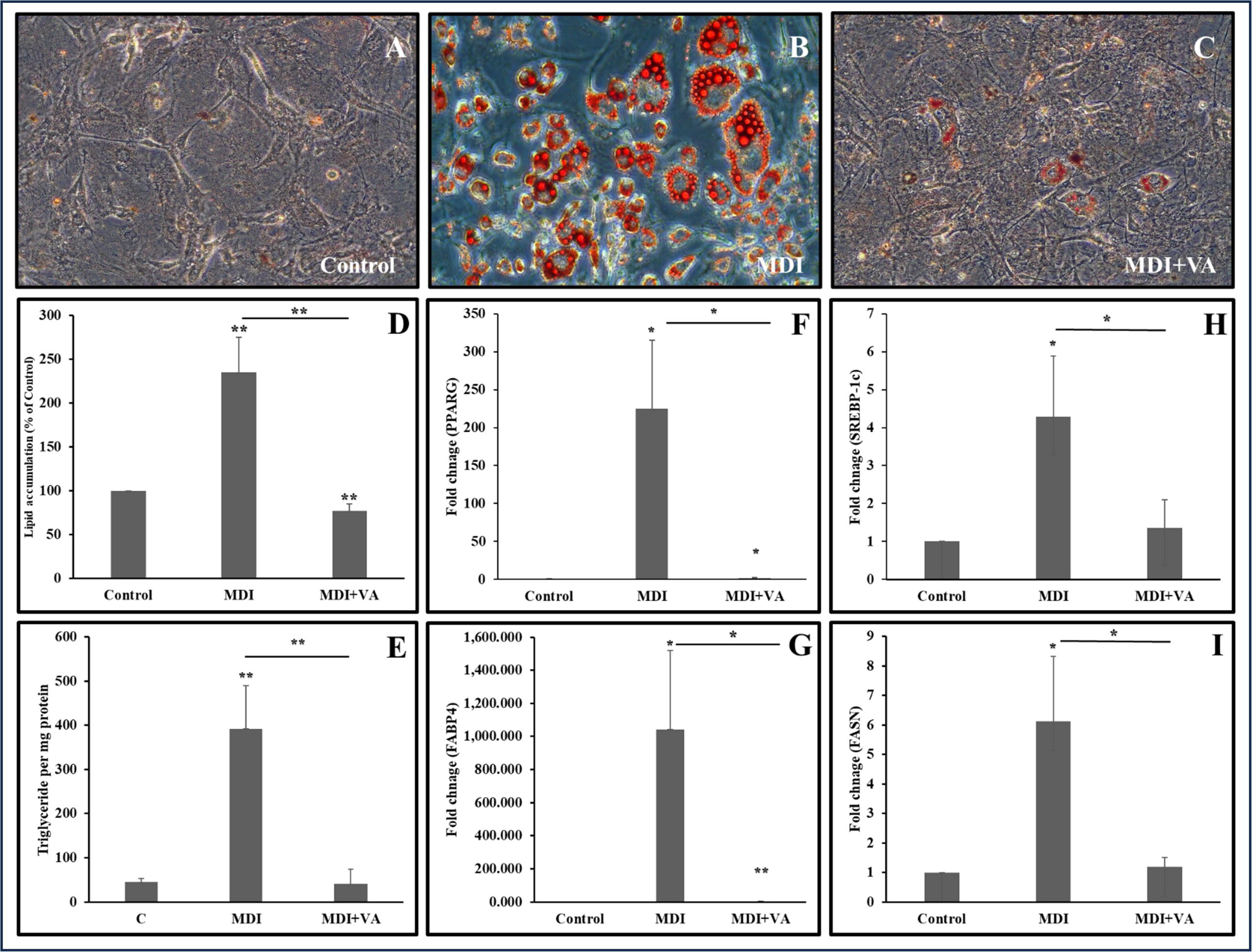
Anti-adipogenic effect of VA on 3T3-L1 cells. Microscopic image of cells showing the Oil-Red-O staining of 3T3-L1 cells (20x). **(A)** Control, **(B)** MDI, **(C)** MDI+VA cells and **(D)** the quantification of oil droplets by Oil-Red-O staining and **(E)** Triglyceride content. **F – I** depicts the qRT-PCR results showing the modulation of adipogenic genes such as **(F)** *Ppar*γ, **(G)** *Fabp4*, **(H)** *Srebp-1c* and **(I)** *Fasn*. Data represented as mean ± SD (p value ≤ ** 0.01, p value ≤ * 0.05).

### 3.4 VA treatment improved the HFD-STZ mediated histopathological anomalies in pancreas, liver, kidney and intestine in experimental animals

Histopathological analysis of key organs revealed the protective effects of VA treatment in HFD-STZ-induced diabetic rats. In the pancreas, VA-treated animals exhibited reduced inflammation and an increase in pancreatic beta islet cells compared to the diabetic group, indicating improved pancreatic morphology. In the liver, HFD-STZ rats displayed moderate inflammation, steatosis, and associated inflammatory responses, while VA-treated groups demonstrated improved hepatic architecture, with reduced steatosis and inflammation. Renal histopathology showed that VA-treated kidneys had reduced inflammation and preserved glomerular and tubular structures, suggesting a protective effect on renal tissues. Intestinal examination revealed that VA treatment mitigated damage to the intestinal lining, reduced inflammation, and prevented edema. In contrast, the HFD-STZ group exhibited extensive epithelial damage, loss of goblet cells, and pronounced inflammatory infiltration. These findings suggest that VA treatment effectively alleviates inflammation and preserves cellular integrity across multiple organs in the context of HFD-STZ-induced diabetes (Fig. 3).

**Fig. 3.**
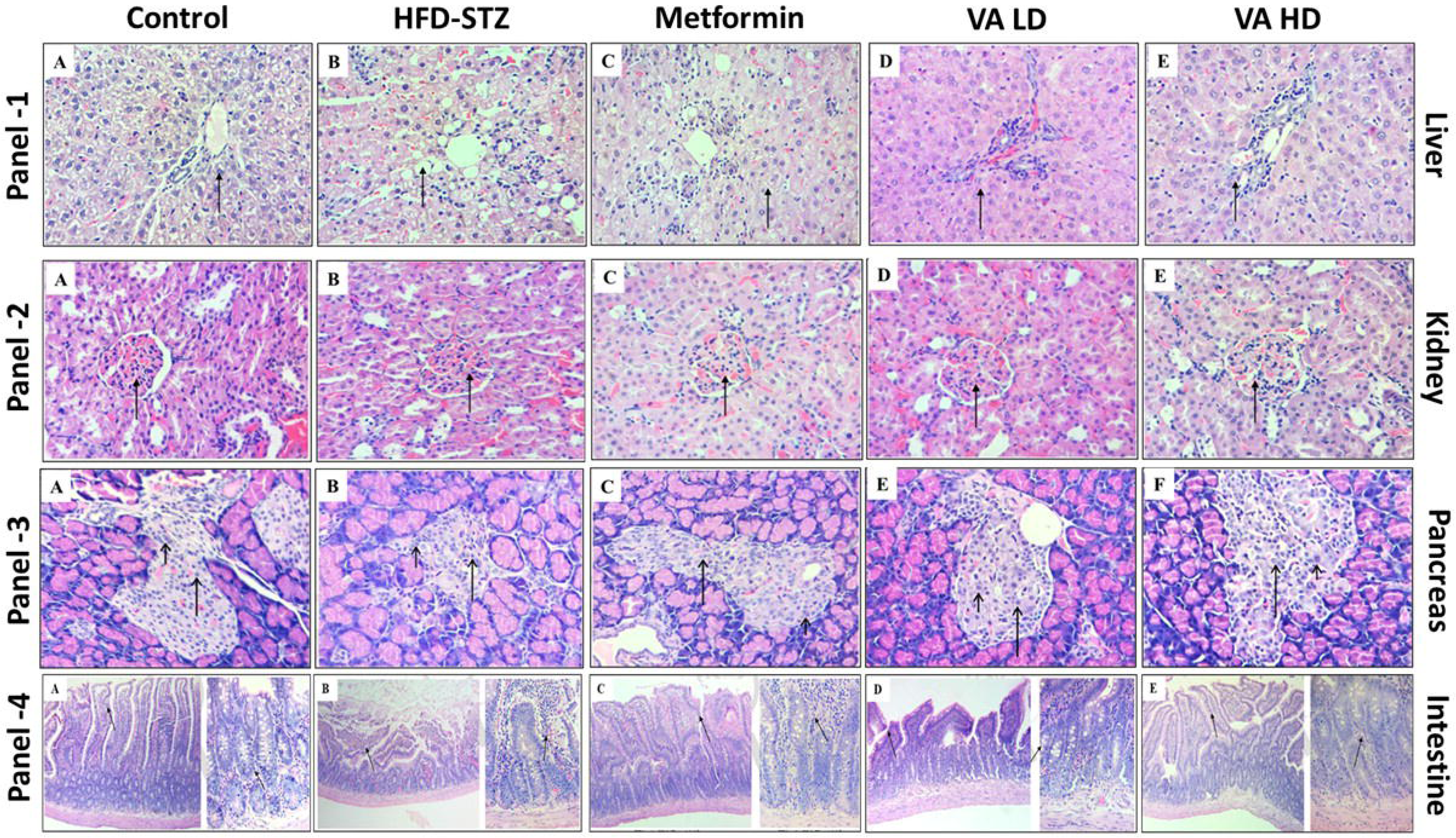
Histopathology of rat tissues. Images showing the histopathology results of Liver, Kidney, Pancreas and Intestine from HFD-STZ fed SD rats treated with metformin, VA LD and VA HD.

### 3.5 VA modulates incretin effect by inhibiting DPP4 enzyme activity, as well as increasing GLP-1 secretion and inhibiting DPP4 expression in GLUTag cells

One of the key players of the incretin effect is the secretion of GLP-1 hormone from enteroendocrine cells. However, as soon as the hormone is secreted by the enteroendocrine cells, the serine protease DPP4 will degrade it and thereby reduce their biological effects. Considering the vital role DPP4 plays in incretin biology, DPP4 inhibitors are considered to be important in diabetes management. In our study, we found that VA inhibits DPP4 enzyme activity in a dose dependent manner with an IC_50_ of 1.22 μg of GAE/ml. At the highest tested concentration of 50μg of GAE/ml, VA inhibited DPP4 activity by 97% and at lowest tested concentration of 6.25μg of GAE/ml VA showed an inhibition of 73% (Fig. 4A).

**Fig. 4.**
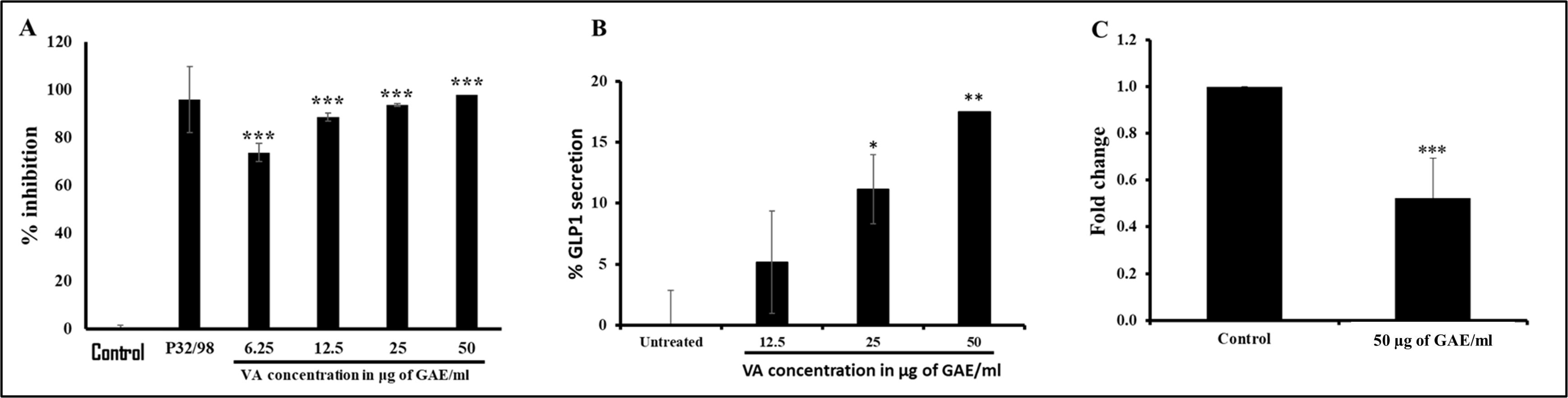
DPP4 inhibition effect of VA and GLP-1 secretion in GLUTag cells. **(A)** Graph shows a concentration-dependent inhibition of DPP4 enzyme. **(B)** Graph shows a concentration dependent increase in GLP1 release from GLUTag cells upon treatment with various concentrations of VA **(C)** Graph shows a downregulation of DPP4 expression at 50 µg of GAE/ml VA. Data represented as mean ± SD (p value ≤ *** 0.001, p value ≤ ** 0.01, p value ≤ * 0.05).

Additionally we used the GLUTag cells, a murine enteroendocrine cell line that expresses the proglucagon gene and secretes the glucagon-like peptides similar to the human system. GLUTag cells treated with VA showed a concentration-dependent increase in GLP-1 secretion which further supports the increased serum GLP-1 levels observed in SD rats (Fig. 4B). In addition to increased GLP-1 secretion, studies have shown that inhibition of DPP4 expression at mRNA level reduces the circulating concentration of DPP4 and it is also a strategy for positively modulating the incretin effect^20^. To study this angle, GLUTag cells were treated with 50µg of GAE/mL of VA and our results showed 1.5-fold reduction in DPP4 expression compared to the control (Fig. 4C). Taken together, these results support the possible incretin modulation effect of VA in exerting its anti-diabetic effects.

### 3.6 Molecular docking and molecular dynamics simulation studies identified key phytochemicals in VA that inhibit DPP4 activity

Considering the importance of DPP4 in diabetes management and the observed DPP4 inhibition effect of VA, we have employed computational biology methods to identify the phytochemicals in VA that could potentially inhibit DPP4 enzyme action and analysed their molecular dynamics. A total of 248 phytochemicals were identified from the 16 constituent plants in VA and subjected to molecular docking with DPP4 protein and MM/GBSA studies using Vildagliptin as a standard DPP4 inhibitor. The top 20 compounds, ranked based on their docking scores and MM/GBSA values, are presented in Table 6. Among these, Chebulinic acid and Chebulagic acid emerged as the most promising candidates and were thus selected for detailed molecular dynamics and simulation studies. While other compounds such as 1,2,3,4,6-Penta-O-galloyl-α-D-glucopyranose (2nd rank), Terflavin A (3rd rank), Pentagalloyl Glucose (4th rank), and Terchebin (5th rank) also exhibited favorable docking and MM/GBSA scores, Chebulinic acid and Chebulagic acid were prioritized due to their distinctive interaction profiles and potential as lead compounds. 1,2,3,4,6-Penta-O-galloyl-a-D-glucopyranose, Terflavin A and Pentagalloyl Glucose were found to be not showing stable interactions with the DPP4 whereas Terchebin was already studied for its molecular dynamics and simulation with DPP4 in our previous study on another formulation *Nishakathakadi Kashayam*.^19^ The docking and MM/GBSA scores for the standard Vildagliptin (−3.899 kcal/mol and −26.7 kcal/mol) was also taken from the previous study.^19^ The MM/GBSA scores of Chebulinic acid and Chebulagic acid were found to be, −64.61, and −46.51 kcal/mol whereas the docking scores were, −10.631 and −9.322 kcal/mol respectively (Table 6).

**Table 6.**
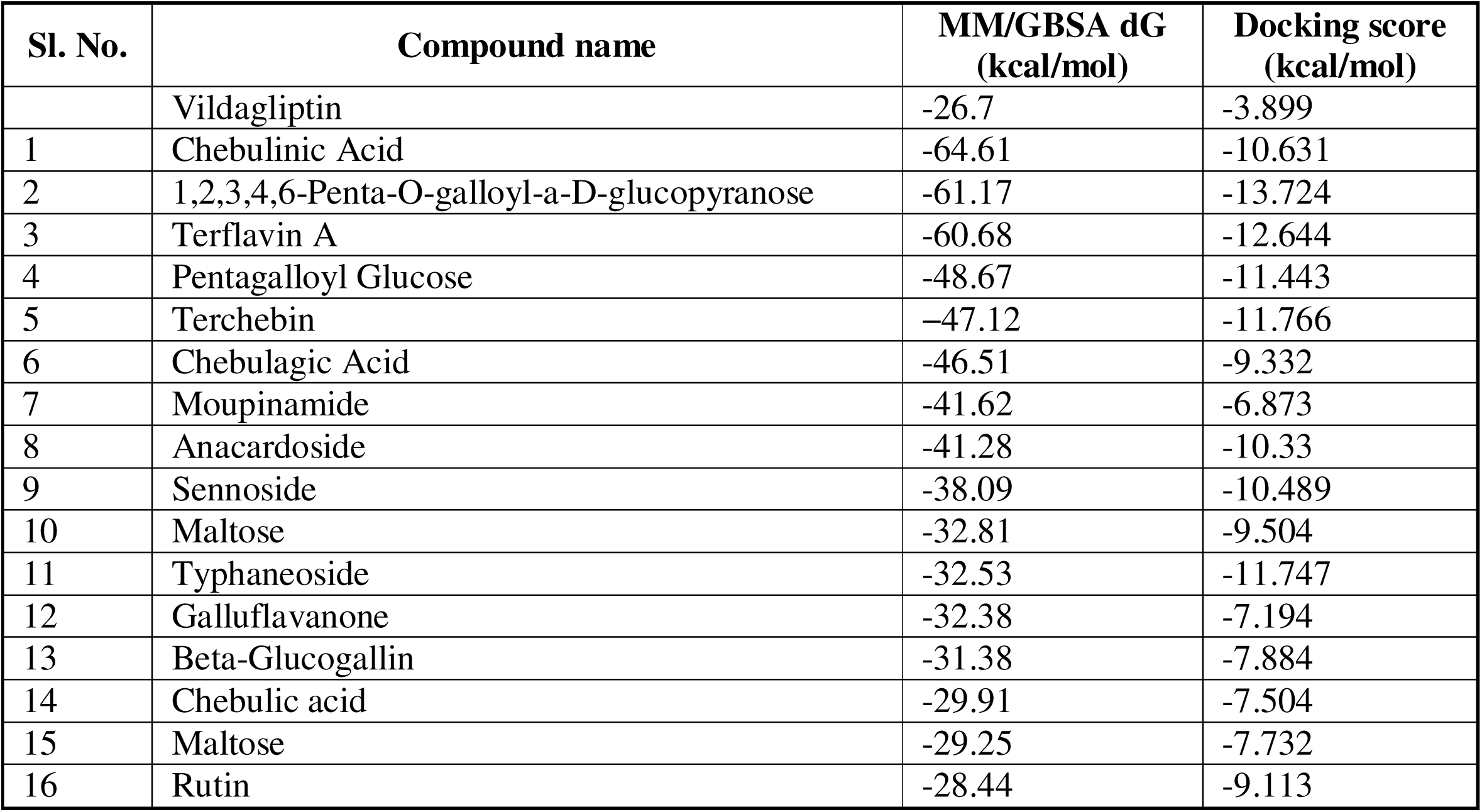
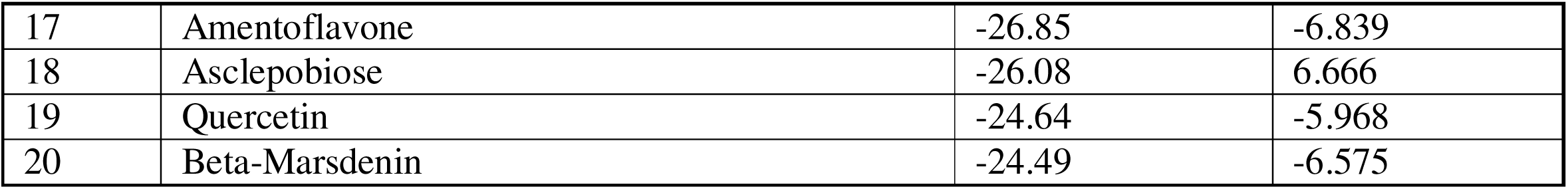
The top-ranked 20 compounds, based on their docking and MM/GBSA scores against DPP4.

MD is a powerful computer simulation technique that is now being used in the field of computer-aided drug discovery research to deepen the understanding of how the protein-ligand complex behaves in a dynamic environment at the atomic level over a user-specified time period. A 100ns MD simulation analysis of the DPP4-ligand docked complex was performed and the RMSD profiles of the protein and ligands during the 100ns simulations are analysed. In the DPP4-Chebulinic acid complex (Fig. 5C), the protein Cα RMSD stabilized around 1.8 Å during the last 50ns, indicating overall protein stability. However, the ligand RMSD showed a marked increase after ∼40ns, suggesting a shift in ligand binding. In contrast, the DPP4-Chebulagic acid complex (Fig. 5D) exhibited stable protein RMSD around 1.6 Å and consistently high but stable ligand RMSD throughout the 100ns simulation, indicating a more stable and persistent ligand–protein interaction. Chebulinic acid was found to directly interact with key residues, including Glu205, Lys554, Asn562, Trp629, and His740 (Fig. 5A), while chebulagic acid showed direct interactions with Glu205, Pro550, Arg560, Ser630, Tyr547, Gly741, and His740 (Fig. 5B) (Supplementary Fig. 1). Virtual screening results showed that both chebulinic acid and chebulagic acid interacted with key residues involved in Vildagliptin binding, such as Glu205 and His740.^19^ These interactions are stabilized by prominent hydrogen bonds, water bridges and π-π stacking that significantly contribute for a stable ligand-protein interaction.

**Fig. 5.**
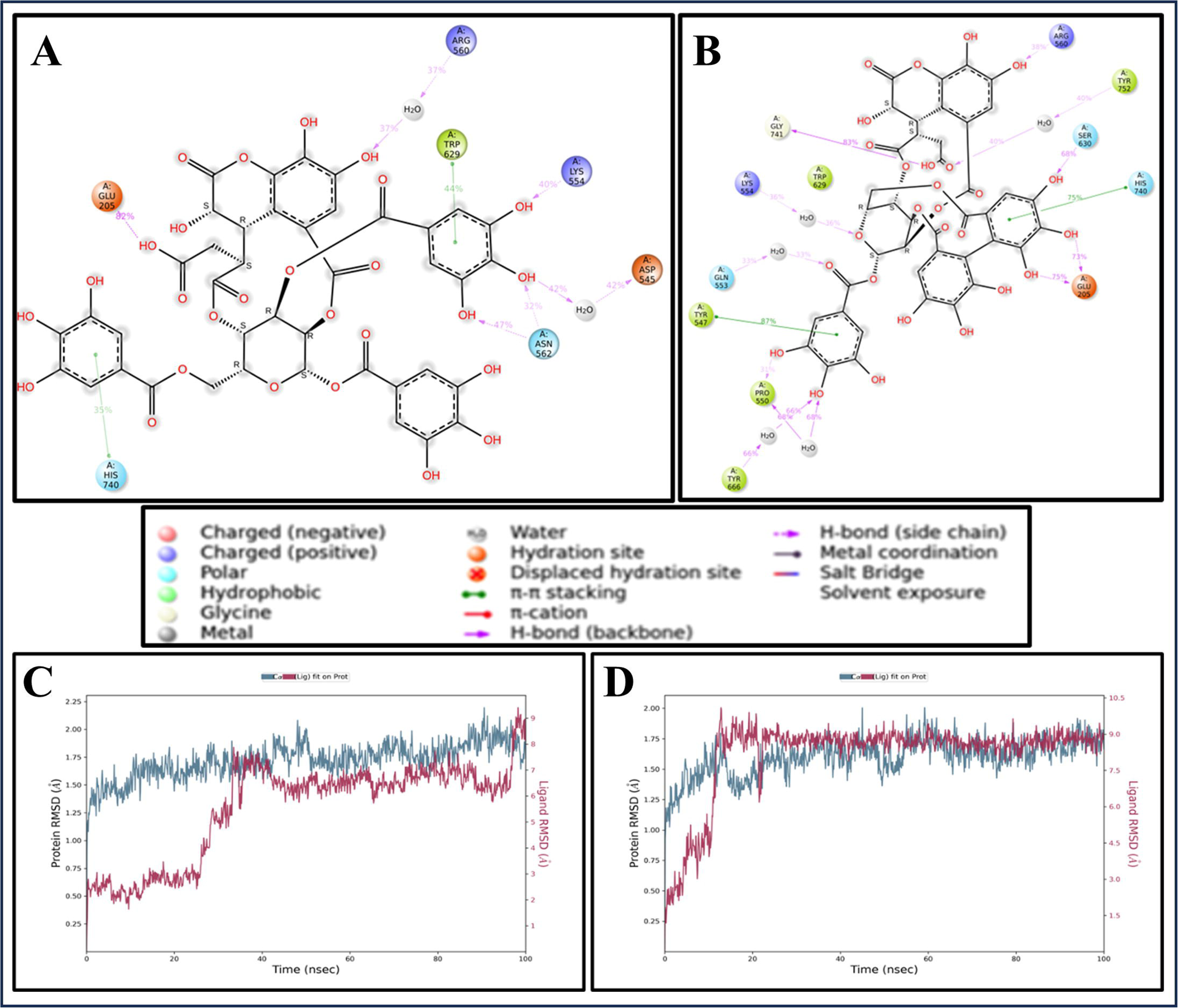
Interaction of DPP4 with various ligands and Protein-ligand RMSD plot. **2D** interaction diagrams of DPP4 with **(A)** Chebulinic acid and **(B)** Chebulagic acid. RMSD plots of protein-ligand complexes, **(C)** DPP4-Chebulinic acid and **(D)** DPP4-Chebulagic acid.

## 4. Discussion

As per *Ayurveda* epistemology, the polyherbal formulation *Varanadi Kashayam* (VA) is prescribed for correcting the “*Agni*” imbalance and “*Dosha*” dysregulations, that are often associated with the pathophysiology of diseases like diabetes, obesity, chronic liver diseases, abdominal bloating, chronic arthritis etc.^9^ *Ashtanga Hridaya* and *Susrutha samhitha*, two classical texts of *Ayurveda*, describes a category of plants as ‘*Varanadi gana*’ having therapeutic indications for diseases arising from impaired *Agni* and vitiated *Dosha* functions as well as dyslipidemia.^9,11^ VA comprises 16 herbs belonging to this *Varanadi gana*, many of which are extensively studied for their biological activities related to metabolic diseases and other disease conditions.^25–29^ While understanding the mechanism of ingredient plants is essential, assessing the formulation as a whole is equally important. This is because the medicinal chemistry and network pharmacology of a formulation like VA would be different from the sum of their parts owing to combinatorial effects and methods of processing. There are three important studies on VA by Chinju *et al*, where the formulation is studied in its *Kashaya* form reporting its anti-inflammatory and anti-obesogenic effects using both *in-vitro* and *in-vivo* model systems.^23,24,30,31^ These findings align with the Ayurvedic claims of therapeutic benefits of *Varanadi Gana*.

Most of the ingredient plants of VA are well-studied for their medicinal properties in diabetes and other metabolic diseases. ^32, 33, 34^ These herbs exhibit key anti-diabetic action such as alpha-glucosidase inhibition, protein glycation, DPP4 inhibition and enhancement of GLP-1 secretion. Specifically, *Plumbago zeylanica, Aegle marmelos, Pongamia pinnata, Moringa oleifera and Terminalia chebula* are shown to DPP4 enzyme action while *Asparagus racemosus, Aegle marmelos, Pongamia pinnata* are reported to increase GLP-1 secretion in various models. ^19,35,36,37^ Similarly *Terminalia chebula* and *Moringa oleifera* are studied for their potential to improve impaired glucose tolerance. ^38,39^ Collectively, the anti-hyperglycemic effects of VA may stem from a multi-component multi-targeted synergistic mechanism involving these herbs. ^40–42^

Our current study evaluates the anti-diabetic effects of VA with a particular focus on its ability to modulate the incretin effect. In Ayurvedic parlance, restoration of *Agni* - the digestive and metabolic fire - is central to treating metabolic diseases. *Agni Mandya* (impaired digestion and metabolic functions) is considered to be the primary cause of various metabolic diseases, including diabetes and obesity. Consequently, Ayurvedic treatment strategies emphasize the correction of *Agni Mandya* through herbal drug interventions. Both *Ayurveda* and modern biology epistemology converges and recognizes GIT as a central regulator of systemic glucose homeostasis. Building on this integrative framework, we have previously correlated Ayurvedic *Agni*-based diabetes management with gastrointestinal mediated glucose disposal (GIGD), which involves a complex network of gut derived molecular regulators of glucose metabolism. ^8^ Among these, incretin hormones play a pivotal role in enhancing insulin responsiveness and modulating several other biochemical pathways. Therapeutic strategies such as GLP1 agonists and DPP4 inhibitors, which enhances this incretin effect are cornerstones in the management of type 2 diabetes and obesity.^43^ Recent advancements include multi-agonist drugs targeting multiple gut hormones, providing a more comprehensive approach to metabolic disease management.^44^ Despite the availability of approved incretin modulators, the search for safer, more efficacious and more systemically acting molecules/formulations is ongoing. This is the premise in which our study becomes significant by exploring the possible incretin modulatory effect of VA using *in vitro* and *in vivo* model systems.

The *in vivo* high-fat diet low dose streptozotocin rat model employed in our study effectively replicates essential features of T2D such as obesity, insulin resistance and persistent hyperglycemia, while minimizing animal mortality.^45^ This chronic-induction model is particularly suitable for mimicking the multi-system pathophysiology of T2D and therefore exploring the multi-system targeted mode of action of VA that is described in *Ayurveda*. An acute disease model like alloxan or high-dose of STZ might not be relevant to capture the *Medohara* (anti-obesity), *Agni Vardhaka* (gut-modulatory) and *Doshahara* (balancing body humors) properties attributed to VA. Also, the experimental model is well suited for studying the incretin biology due to its high-fat diet induced biochemistry and metabolic profile.

In our study, induction with HFD-STZ showed a significant increase in the fasting blood glucose level and impaired oral glucose tolerance, both hallmark indicators of insulin resistance and type-2 diabetes. A 30-days treatment of the diabetic groups with VA significantly reduced the fasting blood glucose and improved the OGTT compared to untreated diabetic control. OGTT is a crucial evaluation that reflects the patient’s ability to effectively use and store glucose within 2-3 hrs after a meal. It is widely employed to diagnose insulin resistance, pre-diabetes and diabetes.^46^ A crucial contributor to this postprandial glucose regulation is the incretin effect - the enhanced insulin response elicited by nutrient ingestion. It is well established that the incretin effect is severely compromised in T2D.^47^ In our study, we found that HFD-STZ induction reduced the serum levels of GLP-1, one of the marker hormones of incretin effect, and VA treatment restored it suggesting its putative incretin modulatory effect. This result partly explains the positive modulation of FBS and OGTT observed in VA treated animals.

In humans, incretin hormones are secreted by the enteroendocrine cells lining the GI tract in response to glucose and other nutrients. The key incretin hormone, GLP-1, is secreted by the L cells located primarily in the ileum and large intestine. To confirm the increased serum GLP-1 levels observed in experimental animals treated with VA, we utilized GLUTag cells, a murine enteroendocrine cells generated by Prof. Drucker’s lab.^48^ GLUTag cells are widely recognized as gold-standard for *in vitro* studies related to incretin biology due to their robust expression of proglucagon gene and secretion of incretin hormones, especially GLP-1. In humans, shortly after the release, GLP-1 and other incretin hormones are rapidly degraded by DPP4, an evolutionarily conserved serine protease, which limits their biological action. Given the critical physiological and therapeutic roles of incretin hormones like GLP-1, strategies that enhance GLP-1 secretion, utilize GLP-1 analogues and targeted inhibition of the DPP4 enzyme remain important areas of research and drug development. Additionally, studies have also shown that the downregulation of DPP4 expression, particularly in intestinal cells, represent another viable approach to increase the incretin effect and improve the glucose metabolism. Corroborating with our *in-vivo* results, VA treatment increased GLP-1 secretion from GLUTag cells and inhibited DPP4 enzyme activity. We observed that VA treatment increases GLP-1 secretion in GLUTag cells as well as inhibits DPP4 enzyme action. Also, the expression of DPP4 was downregulated in VA treated intestinal cells.

Given the significant DPP4 inhibition observed, we explored the potential bioactive compounds responsible for this activity. Data mining followed by molecular docking studies showed several bioactive compounds that can potentially interact with DPP4, three compounds viz. chebulinic acid, terchebin and chebulagic acid came as top candidates with satisfying molecular dynamic simulation results. Chebulinic acid, an ellagitannin, and chebulagic acid, a benzopyran tannin, are reported from *Terminalia chebula*, and are known for their anti-inflammatory and antioxidant properties. Recently, Yan et. al., 2024 showed through computational biology tools that both chebulagic acid and chebulinic acid exerts possible antidiabetic effect by interacting with protein tyrosine phosphatase 1B (a crucial enzyme that regulate insulin and leptin signaling) and alpha glucosidase (a digestive enzyme important in glucose metabolism) by displaying substantial interactions with important amino acid residues of the enzyme.^49^ Additionally, chebulagic acid and chebulinic acid are also found to exert various other biological activities like anti-bacterial, anti-cancer, anti-hypertensive and anti-oxidant activities. Terchebin is another compound reported from various plants including Terminalia chebula. Although much studies are not reported on the bioactivities of terchebin, a recent study from our lab identified terchebin in another formulation called *Nishakathakadi Kashayam* and has shown its DPP4 inhibition activity.^19^

Another important Ayurvedic claim of this formulation is its potential hypocholesterolemic and anti-lipidemic effects that are essential in restoring a systemic glucose homeostasis in the body. Accumulation of triglycerides is one of the conditions found to have a bidirectional relation (cause and effect) in impaired glucose metabolism and associated disease conditions. ^50^ Our results with experimental animals as well as the fibroblast cell model indicate that VA can reduce the triglyceride level both systemically and in the cell model system. Both plasma triglyceride and plasma cholesterol levels were reduced in VA-treated groups demonstrating a hypocholesterolemic and anti-lipidemic effects of the formulation, which are in line with the previously reported anti-obesity effects of the formulation.^23^ Also, these results support the *Ayurveda* indications of these ‘*kashayas*’ (decoctions) as substances that are correcting the irregular metabolism and improve digestive functions to reduce the lipids or ‘*meda*’, in plasma levels.

## 5. Conclusions

In summary, our study used a combined approach of *in vitro*, *in vivo* and *in silico* methods to delineate the possible anti-diabetic action of VA and to partly correlate it with Ayurveda pharmacology. Our observations suggest that VA exert its anti-diabetic activity through a gut-centred multi-targeted mode of action prominently involving incretin modulation. Additionally the identification of bioactive compounds like chebulagic acid, chebulinic acid and terchebin provides a strong basis for future drug development.

## List of Abbreviations

T2D: Type 2 Diabetes
GLP-1: Glucagon like peptide - 1
DPP4: Dipeptidyl Peptidase-4
GIT: Gastrointestinal Tract
NAFLD: Non-Alcoholic Fatty Liver Diseases
GAE/mL: Gallic Acid Equivalent per milliliter
DMEM: Dulbecco’s Modified Eagle’s Medium
FBS: Fetal Bovine Serum
PBST: Phosphate buffered saline with tween-20
IBMX: 3-isobutyl-1-methylxanthine
*Fabp4*: Fatty acid binding protein 4
*Fasn*: Fatty Acid Synthase
*Gapdh*: Glyceraldehyde 3-phosphate dehydrogenase
*Ppar*γ: Peroxisome proliferator-activated receptor gamma
*Srebp-1c*: Sterol Regulatory Element Binding Protein 1c
FP: Forward primer
RP: Reverse primer
ELISA: Enzyme Linked Immunosorbent Assay
SD Rats: Sprague Dawley rats
HFD-STZ: high fat diet and streptozotocin
HD: High Dose
LD: Low Dose
OGTT: Oral glucose tolerance test
BW: Body weight
ANOVA: Analysis of Variance
AUC: Area under curve
IMPPAT: Indian Medicinal Plants, Phytochemistry and Therapeutics
MD Simulation: Molecular Dynamics simulation
RMSD: Root mean square deviation.

## Acknowledgement

The authors acknowledge the financial support received by Anjana from Tata Education & Development Trust as well as Rural India Support Trust (RIST); Dr Daniel Drucker for giving his permission to use GLUTag cells and Dr Tohru Hira for supplying the cells; financial support received by Sania Kouser (ICMR-SRF) from Indian Council of Medical Research (ICMR (3/1/2(13)/OBS/2022-NCD-II)); the financial support for a Research Fellow from TDU, Bangalore; JM Financials for supporting the research project through TDU; and SASTRA Deemed to be University and DBT (grant no. BT/PR40144/BTIS/137/46/2022) for Schrodinger software (https://www.schrodinger.com/) support. The authors thank Sthitaprajna Sahoo and Abhijnan Chakraborty for contributing to the docking part.

## Author Contributions

CNVP, SKK and SS conceptualized the work; experiments were performed by AT, SK, AVB, PGP and SR; data validation was done by AT, SK, CNVP, SKK, SS and SJ and MJM.; resources for the experiments were arranged by CNVP, SS, SJ and MJM.; Data curation and original manuscript writing was done by AT, SK, AVB, SS and CNVP and review of manuscript was done by CNVP, SJ, SKK and SS. All authors have read and agreed to the published version of the manuscript.

---

**Supplementary Fig. 1.** Prominent hydrogen, hydrophobic, ionic bonds, and water bridges that significantly contribute to a stable ligand protein interaction. **(A)** DPP4-Chebulinic acid and **(B)** DPP4-Chebulagic acid.

## References

1. King A, Miller EM. Glucagon-Like Peptide 1 Receptor Agonists Have the Potential to Revolutionize the Attainment of Target A1C Levels in Type 2 Diabetes—So Why Is Their Uptake So Low? Clinical Diabetes. 2023;41(2):226–238. doi:10.2337/cd22-0027

2. Association AD. 1. Improving Care and Promoting Health in Populations: *Standards of Medical Care in Diabetes—*2021. Diabetes Care. 2021;44(Supplement_1):S7–S14. doi:10.2337/dc21-S001

3. Drucker DJ. Mechanisms of Action and Therapeutic Application of Glucagon-like Peptide-1. Cell Metab. 2018;27(4):740–756. doi:10.1016/j.cmet.2018.03.001

4. Deacon CF. Peptide degradation and the role of DPP-4 inhibitors in the treatment of type 2 diabetes. Peptides (NY). 2018;100:150–157. doi:10.1016/j.peptides.2017.10.011

5. Guo W, Xu Z, Zou H, et al. Discovery of ecnoglutide – A novel, long-acting, cAMP-biased glucagon-like peptide-1 (GLP-1) analog. Mol Metab. 2023;75:101762. doi:10.1016/j.molmet.2023.101762

6. Mathur V, Alam O, Siddiqui N, et al. Insight into Structure Activity Relationship of DPP-4 Inhibitors for Development of Antidiabetic Agents. Molecules. 2023;28(15):5860. doi:10.3390/molecules28155860

7. Michaelidou M, Pappachan JM, Jeeyavudeen MS. Management of diabesity: Current concepts. World J Diabetes. 2023;14(4):396–411. doi:10.4239/wjd.v14.i4.396

8. Thottapillil A, Kouser S, Kukkupuni SK, Vishnuprasad CN. An ‘Ayurveda-Biology’ platform for integrative diabetes management. J Ethnopharmacol. 2021;268:113575. doi:10.1016/j.jep.2020.113575

9. Shastri A. Sushruta Samhita, Ayurveda-Tattva-Samdipika Commentary.; 2003.

10. Murthy KRS. Astanga Hrdayam. CHOWKHAMBA.; 2022.

11. Singh RH. Ashtanga Hridayam. a.; 2013.

12. Dwarakanath C. The Fundamental Principles of Ayurveda.; 2003.

13. Ainsworth EA, Gillespie KM. Estimation of total phenolic content and other oxidation substrates in plant tissues using Folin–Ciocalteu reagent. Nat Protoc. 2007;2(4):875–877. doi:10.1038/nprot.2007.102

14. Butala MA, Kukkupuni SK, Vishnuprasad CN. Ayurvedic anti-diabetic formulation Lodhrasavam inhibits alpha-amylase, alpha-glucosidase and suppresses adipogenic activity in vitro. J Ayurveda Integr Med. 2017;8(3):145–151. doi:10.1016/j.jaim.2017.03.005

15. Kifle ZS, Belayneh YM. Antidiabetic and Anti-hyperlipidemic Effects of the Crude Hydromethanol Extract of *Hagenia abyssinica* (Rosaceae) Leaves in Streptozotocin-Induced Diabetic Mice. Diabetes Metab Syndr Obes. 2020;Volume 13:4085–4094. doi:10.2147/DMSO.S279475

16. Ghosh MN. Fundamentals of experimental pharmacology. 6th Ed Kolkata:Hilton and company. Published online 2015.

17. Ghasemi & Jeddi S. A. Streptozotocin as a tool for induction of rat models of diabetes: a practical guide. EXCLI J. 2023;22(274–294).

18. Mohanraj K, Karthikeyan BS, Vivek-Ananth RP, et al. IMPPAT: A curated database of Indian Medicinal Plants, Phytochemistry And Therapeutics. Sci Rep. 2018;8(1):4329. doi:10.1038/s41598-018-22631-z

19. Thottappillil A, Sahoo S, Chakraborty A, et al. *In vitro* and *in silico* analysis proving DPP4 inhibition and diabetes-associated gene network modulation by a polyherbal formulation: *Nisakathakadi Kashaya*. J Biomol Struct Dyn. 2024;42(24):13588–13602. doi:10.1080/07391102.2023.2276880

20. Lalitha N, Sadashivaiah B, Ramaprasad TR, Singh SA. Anti-hyperglycemic activity of myricetin, through inhibition of DPP-4 and enhanced GLP-1 levels, is attenuated by co-ingestion with lectin-rich protein. PLoS One. 2020;15(4):e0231543. doi:10.1371/journal.pone.0231543

21. Bhat GA, Khan HA, Alhomida AS, Sharma P, Singh R, Paray BA. GLP-I secretion in healthy and diabetic Wistar rats in response to aqueous extract of Momordica charantia. BMC Complement Altern Med. 2018;18(1):162. doi:10.1186/s12906-018-2227-4

22. Furman BL. StreptozotocinLInduced Diabetic Models in Mice and Rats. Curr Protoc. 2021;1(4). doi:10.1002/cpz1.78

23. Chinchu JU, Mohan MC, Prakash Kumar B. Anti-obesity and lipid lowering effects of Varanadi kashayam (decoction) on high fat diet induced obese rats. Obes Med. 2020;17:100170. doi:10.1016/j.obmed.2019.100170

24. J.U C, Mohan MC, B PK. Downregulation of adipogenic genes in 3T3-L1 Pre adipocytes-a possible mechanism of anti-obesity activity of herbal decoction Varanadi Kashayam. J Herb Med. 2020;19:100309. doi:10.1016/j.hermed.2019.100309

25. Akanji MA, Olukolu SO, Kazeem MI. Leaf Extracts of *Aerva lanata* Inhibit the Activities of Type 2 DiabetesLRelated Enzymes and Possess Antioxidant Properties. Oxid Med Cell Longev. 2018;2018(1). doi:10.1155/2018/3439048

26. Vadivelan R, Gopala Krishnan R, Kannan R. Antidiabetic potential of Asparagus racemosus Willd leaf extracts through inhibition of α-amylase and α-glucosidase. J Tradit Complement Med. 2019;9(1):1–4. doi:10.1016/j.jtcme.2017.10.004

27. Kesari AN, Gupta RK, Singh SK, Diwakar S, Watal G. Hypoglycemic and antihyperglycemic activity of Aegle marmelos seed extract in normal and diabetic rats. J Ethnopharmacol. 2006;107(3):374–379. doi:10.1016/j.jep.2006.03.042

28. Muhammad HI, Asmawi MZ, Khan NAK. A review on promising phytochemical, nutritional and glycemic control studies on Moringa oleifera Lam. in tropical and sub-tropical regions. Asian Pac J Trop Biomed. 2016;6(10):896–902. doi:10.1016/j.apjtb.2016.08.006

29. Murali Y, Anand P, Tandon V, Singh R, Chandra R, Murthy P. Long-term Effects of *Terminalia chebula* Retz. on Hyperglycemia and Associated Hyperlipidemia, Tissue Glycogen Content and in Vitro Release of Insulin in Streptozotocin Induced Diabetic Rats. Experimental and Clinical Endocrinology & Diabetes. 2007;115(10):641–646. doi:10.1055/s-2007-982500

30. Chinchu JU, Mohan M, Devi SJR, Kumar Bp. Evaluation of anti-inflammatory effect of Varanadi Kashayam (decoction) in THP-1-derived macrophages. AYU (An international quarterly journal of research in Ayurveda). 2018;39(4):243. doi:10.4103/ayu.AYU_53_18

31. Chinchu JU, Mohan MC, Prakash Kumar B. Attenuation of obesity related inflammation in RAW 264.7 macrophages and 3T3-L1 adipocytes by varanadi kashayam and identification of potential bioactive molecules by UHPLC-Q-Orbitrap HRMS. Arch Physiol Biochem. 2023;129(4):879–892. doi:10.1080/13813455.2021.1877309

32. Goyal M, Nagori BP, Pareek A, Sasmal D. Aerva lanata: A review on phytochemistry and pharmacological aspects. Pharmacogn Rev. 2011;5(10):195. doi:10.4103/0973-7847.91120

33. Bag A, Bhattacharyya SK, Chattopadhyay RR. The development of Terminalia chebula Retz. (Combretaceae) in clinical research. Asian Pac J Trop Biomed. 2013;3(3):244–252. doi:10.1016/S2221-1691(13)60059-3

34. Shafi KM, Sajeevan RS, Kouser S, Vishnuprasad CN, Sowdhamini R. Transcriptome profiling of two Moringa species and insights into their antihyperglycemic activity. BMC Plant Biol. 2022;22(1):561. doi:10.1186/s12870-022-03938-6

35. Salehi B, Ata A, V. Anil Kumar N, et al. Antidiabetic Potential of Medicinal Plants and Their Active Components. Biomolecules. 2019;9(10):551. doi:10.3390/biom9100551

36. Abiola JO, Oluyemi AA, Idowu OT, et al. Potential Role of Phytochemicals as Glucagon-like Peptide 1 Receptor (GLP-1R) Agonists in the Treatment of Diabetes Mellitus. Pharmaceuticals. 2024;17(6):736. doi:10.3390/ph17060736

37. Dinda B, Dinda M. Natural Products, a Potential Source of New Drugs Discovery to Combat Obesity and Diabetes: Their Efficacy and Multi-targets Actions in Treatment of These Diseases. Natural Products in Obesity and Diabetes. Published online 2022:101–275. doi:10.1007/978-3-030-92196-5_4

38. Agrawal OD, Kulkarni YA. Treatment with Terminalia chebula Extract Reduces Insulin Resistance, Hyperglycemia and Improves SIRT1 Expression in Type 2 Diabetic Rats. Life. 2023;13(5):1168. doi:10.3390/life13051168

39. Ndong M, Uehara M, Katsumata S ichi, Suzuki K. Effects of Oral Administration of Moringa oleifera Lam on Glucose Tolerance in Goto-Kakizaki and Wistar Rats. J Clin Biochem Nutr. 2007;40(3):229–233. doi:10.3164/jcbn.40.229

40. Agrawal R, Sethiya NK, Mishra SH. Antidiabetic activity of alkaloids of *Aerva lanata* roots on streptozotocin-nicotinamide induced type-II diabetes in rats. Pharm Biol. 2013;51(5):635–642. doi:10.3109/13880209.2012.761244

41. Das A, Naveen J, Sreerama YN, Gnanesh Kumar BS, Baskaran V. Low-glycemic foods with wheat, barley and herbs (Terminalia chebula, Terminalia bellerica and Emblica officinalis) inhibit α-amylase, α-glucosidase and DPP-IV activity in high fat and low dose streptozotocin-induced diabetic rat. J Food Sci Technol. 2022;59(6):2177–2188. doi:10.1007/s13197-021-05231-0

42. Aseervatham J, Palanivelu S, Panchanadham S. *Semecarpus anacardium* (Bhallataka) Alters the Glucose Metabolism and Energy Production in Diabetic Rats. Evidence-Based Complementary and Alternative Medicine. 2011;2011(1). doi:10.1155/2011/142978

43. Gimeno RE, Briere DA, Seeley RJ. Leveraging the Gut to Treat Metabolic Disease. Cell Metab. 2020;31(4):679–698. doi:10.1016/j.cmet.2020.02.014

44. Gutgesell RM, Nogueiras R, Tschöp MH, Müller TD. Dual and Triple Incretin-Based Co-agonists: Novel Therapeutics for Obesity and Diabetes. Diabetes Therapy. 2024;15(5):1069–1084. doi:10.1007/s13300-024-01566-x

45. Srinivasan K, Viswanad B, Asrat L, Kaul CL, Ramarao P. Combination of high-fat diet-fed and low-dose streptozotocin-treated rat: A model for type 2 diabetes and pharmacological screening. Pharmacol Res. 2005;52(4):313–320. doi:10.1016/j.phrs.2005.05.004

46. Carmina E, Stanczyk FZ, Lobo RA. Laboratory Assessment. Yen & Jaffe’s Reproductive Endocrinology. Published online 2014:822–850.e3. doi:10.1016/B978-1-4557-2758-2.00034-2

47. Holst J, Vilsboll T, Deacon C. The incretin system and its role in type 2 diabetes mellitus. Mol Cell Endocrinol. 2009;297(1-2):127–136. doi:10.1016/j.mce.2008.08.012

48. Drucker DJ, Jin T, Asa SL, Young TA, Brubaker PL. Activation of proglucagon gene transcription by protein kinase-A in a novel mouse enteroendocrine cell line. Molecular Endocrinology. 1994;8(12):1646–1655. doi:10.1210/mend.8.12.7535893

49. Yan Y, Abdulla R, Xin X, Akber Aisa H. Revealing the potential hypoglycaemic ingredients of Terminalia chebula Retz. by spectrum–effect relationship combining molecular docking and experimental validation. J Funct Foods. 2024;121:106402. doi:10.1016/j.jff.2024.106402

50. Alexopoulos AS, Qamar A, Hutchins K, Crowley MJ, Batch BC, Guyton JR. Triglycerides: Emerging Targets in Diabetes Care? Review of Moderate Hypertriglyceridemia in Diabetes. Curr Diab Rep. 2019;19(4):13. doi:10.1007/s11892-019-1136-3

