## Supplementary figures and images for "The systemic anti-diabetic effect of polyherbal formulation *Varanadi Kashayam* is mediated through GLP-1 secretion and DPP4 inhibition"

### Supplementary Figure - 1

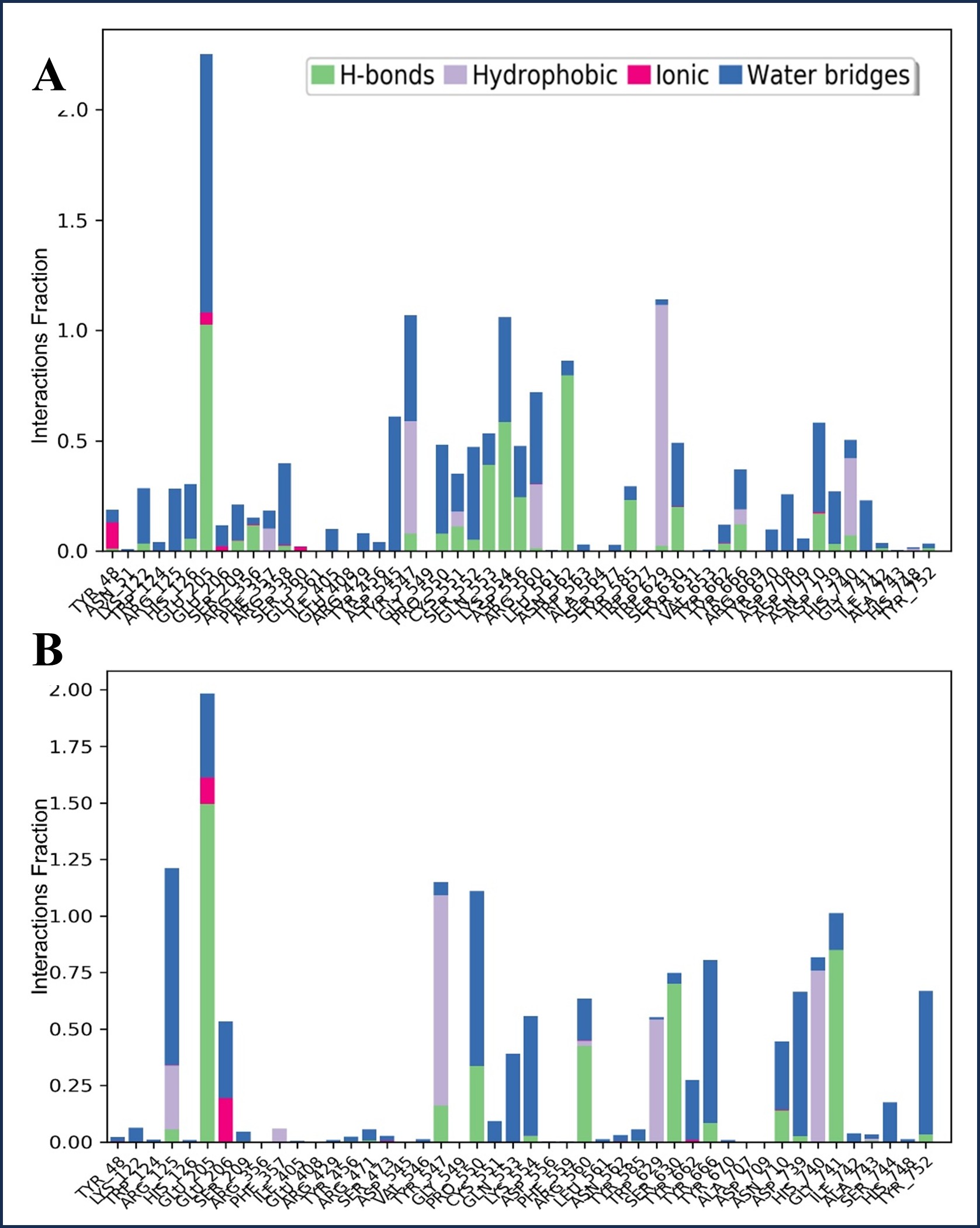
